# Overground walking slip perturbations induce frontal plane motion of the trunk – slips are not just a backwards but also a sideways loss of balance

**DOI:** 10.1101/2023.11.25.568692

**Authors:** Jonathan S. Lee-Confer

**Affiliations:** University of Arizona, Department of Physical Therapy, Tucson, AZ 85721; Musculoskeletal Biomechanics Research Laboratory, University of Southern California, Los Angeles, CA, 90089; Verum Biomechanics, Tucson, AZ 85719

**Keywords:** gait, slip, perturbation, trunk, balance, falls

## Abstract

Slip and fall incidents are a serious health care concern globally. Previous research describes a backwards loss of balance during a slip incident, however hip fractures only occur if individuals fall on their side. Therefore, this study is investigating and quantifying the trunk motion in the sagittal and frontal plane. 13 healthy young participants’ trunk kinematics were analyzed during a slip incident. Peak trunk angle of the trunk in the sagittal and frontal plane were calculated. There was no significant difference between sagittal and frontal plane peak trunk angles suggesting that there is frontal plane motion during an overground slip incident. Our findings suggest research should investigate frontal plane mechanics during a slip incident as there is trunk frontal plane motion which if uncontrolled can result in falling on the femoral neck. Understanding and preventing falls based upon frontal plane mechanics may be more useful for preventing hip fractures from a slip incident. *Lastly, the findings of this study are confirmatory results as the frontal plane trunk motion was quantified and reported in 2008*.

## Introduction

Slip and fall incidents are major health care issue over 300,000 hip fractures occur annually in the United States of America alone [1]. Injuries from falls ranks as the third highest personal health care cost in the United States [2]. Understanding how the human body moves during a slip incident may provide insight on how to regain balance and prevent falls. Research suggests that slip incidences are primarily a sagittal plane perturbation and induce a backwards loss of balance [3]. A slip being described as a backwards loss of balance may be misleading as it is known that hip fractures are far more likely to occur with a sideways loss of balance. As such, the sagittal and frontal characteristics of the body warrant analysis to determine the possibility of there being sideways rotation behavior during an overground slip incident.

The sagittal plane movements of the trunk, arms and legs in response to a slip incident have been reported. Specifically, the mechanics of how the body moves during a slip has been reported to move through the sagittal plane when examining the center of mass [4–6], the upper extremities [5,7–9], the lower extremities [10–12], and the trunk [7,13]. As such, slip research has described a slip incident to result in a backwards loss of balance. The focus of the sagittal plane in a slip incident is a concern as hip fractures do not occur during a backwards loss of balance but only occur when individuals fall to their sides [14–16]. Therefore, it is important to investigate if frontal plane motion is observed during a slip as uncontrolled frontal plane motion would result in a sideways loss of balance and potentially increase risk for a hip fracture.

While most of the slip literature has focused on describing sagittal plane mechanics, there is literature that describes frontal plane mechanics being observed during a slip. Three studies have reported observing frontal plane motion during a slip incident but have not quantified the results [17–19]. Two other studies described frontal plane motion of the arms during a slip incident and showed a photograph of their participant with clear frontal plane trunk motion [20,21]. Nazifi and others (2020) did not quantify shoulder abduction angles during a slip, but a graph in their study shows shoulder abduction angles exceeding that of shoulder flexion angles [9]. Furthermore, Troy and others (2009) conducted a slip study initially starting with 48 individuals and narrowed their subject pool down to 12 by eliminating those whose trunk motion was not directed primarily backwards [7] meaning that other motions, such as frontal or transverse, were present in their participants. Lastly, Smeesters and others (2001) reported that slips mainly result in sideways falls, however their participants were instructed to simulate a faint when the perturbation began and these results may not apply to those who are able to attempt a regaining of balance [22]. These studies, while few, demonstrate that slip incidents may induce frontal plane motion which may be useful for understanding fall mechanics that lead to hip fractures.

As frontal plane trunk motion has not been quantified to date, the purpose of the current study was to characterize the trunk motions in the sagittal and frontal planes during a slip incident. The quantification of peak trunk angles in the sagittal and frontal plane were of interest. We hypothesized the frontal plane trunk flexion would be greater than sagittal plane trunk extension. Information gained from this study will provide insight into how the trunk motion of the human body is induced from a slip incident.

## Methods

### Participants

16 healthy participants between the ages of 21 and 35 participated in this study (8 males and 8 females). Prior to participation, all were informed of the nature of the study, and provided written informed consent as approved by the University of Southern California Health Science Campus Institutional Review Board. After providing informed consent, all participants filled out a medical questionnaire to screen for possible conditions that could jeopardize their safety by participating in the study. Individuals were excluded from participation if they reported any of the following medical conditions: neurological or orthopedic conditions that would affect gait, current muscle strains or joint sprains, recent bone fractures, or previous back injuries.

### Instrumentation

All participant gait trials were performed on a 10-meter walkway within the laboratory. A custom-constructed Nickel-Teflon coated floor tile (California Technical Plating, San Fernando, CA, US) was imbedded into the walkway and camouflaged such that the coloring of the tile matched the non-Teflon tiles. Mineral oil was placed on the tile to reduce the coefficient of friction to induce slipping.

3-D motion analysis was performed using an 11-camera motion analysis system (Qualisys, Gothenburg, Sweden) collected at 150 Hz. 76 reflective markers placed over specific anatomical locations were used to quantify trunk kinematics. To prevent falls during biomechanical testing, a fall-arresting body harness (Miller Model 550-64, Dalloz Fall Protection, Franklin, PA, USA) secured with an 8 mm climbing rope was attached to a low-friction trolley directly above the 10-meter walkway. An Omega S-beam load cell (Omega Engineering Inc., Norwalk, CT, US) was connected the climbing rope and the trolley system to measure the amount of bodyweight displaced on the load cell during the slip perturbation trials. To control for the influence of footwear, all footwear was standardized and participants were fitted with a pair of oxford dress shoes with a standard rubber outer sole (Bates Footwear, Richmond, IN, US).

### Procedures

Prior to testing, an adjustable fall arresting harness was fitted appropriately to each participant. The harness was adjusted so that the hip would not be allowed to drop below a distance equal to 35% of participants’ height [23]. Participants were then instrumented with a full body marker set. Reflective joint markers were placed on the L5S1, Xyphoid Process, and C7, and markers were placed bilaterally on the: second toe, fifth metatarsal head, first metatarsal head, lateral and medial malleolus, lateral and medial epicondyles of the femur, greater trochanter, anterior superior iliac spine, iliac crest, posterior superior iliac spine, acromioclavicular joint, anterior and posterior glenohumeral joint, greater tubercle, lateral and medial epicondyle of the humerus, radial and ulnar styloid processes, and the third metacarpal head. Additionally, a head band fitted with 4 markers was used to track the head, and marker tracking clusters were placed bilaterally on the heel, shank, thigh, upper arm and forearm.

The lighting in the laboratory was dimmed to 3 foot candles prior to the walking trials to assist in concealment of the contaminated tile, but bright enough for participants to navigate the walkway. Participants engaged in multiple practice walking trials to adjust to the harness system and the dimmed lighting conditions until they achieved a consistent walking speed of 1.35-1.5 m/s. All trials (non-slip and slip) monitored gait speed to ensure participants remained within the target speed. To avoid anticipatory gait changes to a potential perturbation, care was taken by slightly dimming the lighting so that participants were unaware of the location of the Teflon tile and which trial the contaminant would be applied [24,25].

Kinematic data were obtained during 4 non-slip walking trials once participants were acclimated to the instrumentation and procedures. Between each trial, participants faced away from the walkway for 60 seconds so that they would be uncertain as to the trial in which a contaminate would be placed on the floor to induce a slip. Loud music was projected throughout the laboratory during each of the 1-minute breaks between trials to act as an additional distraction and avoid the participant detecting contaminant being applied. Mineral was placed on the Teflon tile after obtaining a minimum of 4 non-slip walking trials at the appropriate gait speed. Following the slip trial, participants were asked if they had anticipated the slip trial or if they had observed the contaminant. Any anticipation or observation of the contaminant resulted in the participant being excluded from this study. All participants were slipped on their right foot and were only exposed to 1 slip during the course of the study.

### Data Analysis

For purposes of this study, only data from participants who recovered their balance immediately following the slip were used for analysis. The determinant outcome of the slip, recovery or fall, was determined by the Omega load cell. An outcome was categorized as a fall if the individual displaced more than 30% of their body weight onto the harness system [23].

Kinematic data were filtered using an 2nd order, 6-Hz, low pass Butterworth filter with zero-lag compensation. Fifteen body segments (head, pelvis, thorax, and bilateral feet, shank, thigh, upper arm, forearm, and hand) were created through a custom designed model template using Visual 3D software (C-Motion, Inc., Germantown, MD, USA). The global coordinate system was defined with Y as the anterior-posterior (AP) axis, X as the medio-lateral (ML) axis and Z as the vertical axis. The coordinate system for the thorax were based on the work of Wu and others [26]. Trunk kinematics in the sagittal and frontal plane were exported and analyzed in MATLAB (Mathworks, Natick, MA, USA). Peak sagittal and frontal plane trunk angles after slip initiation were used for analysis. To calculate excursion in each plane, the trunk angle position at the time of heel strike was considered the start position and the final position was the greatest deviation away from the start (Fig. 1 & Fig. 2).

**Figure 1.**
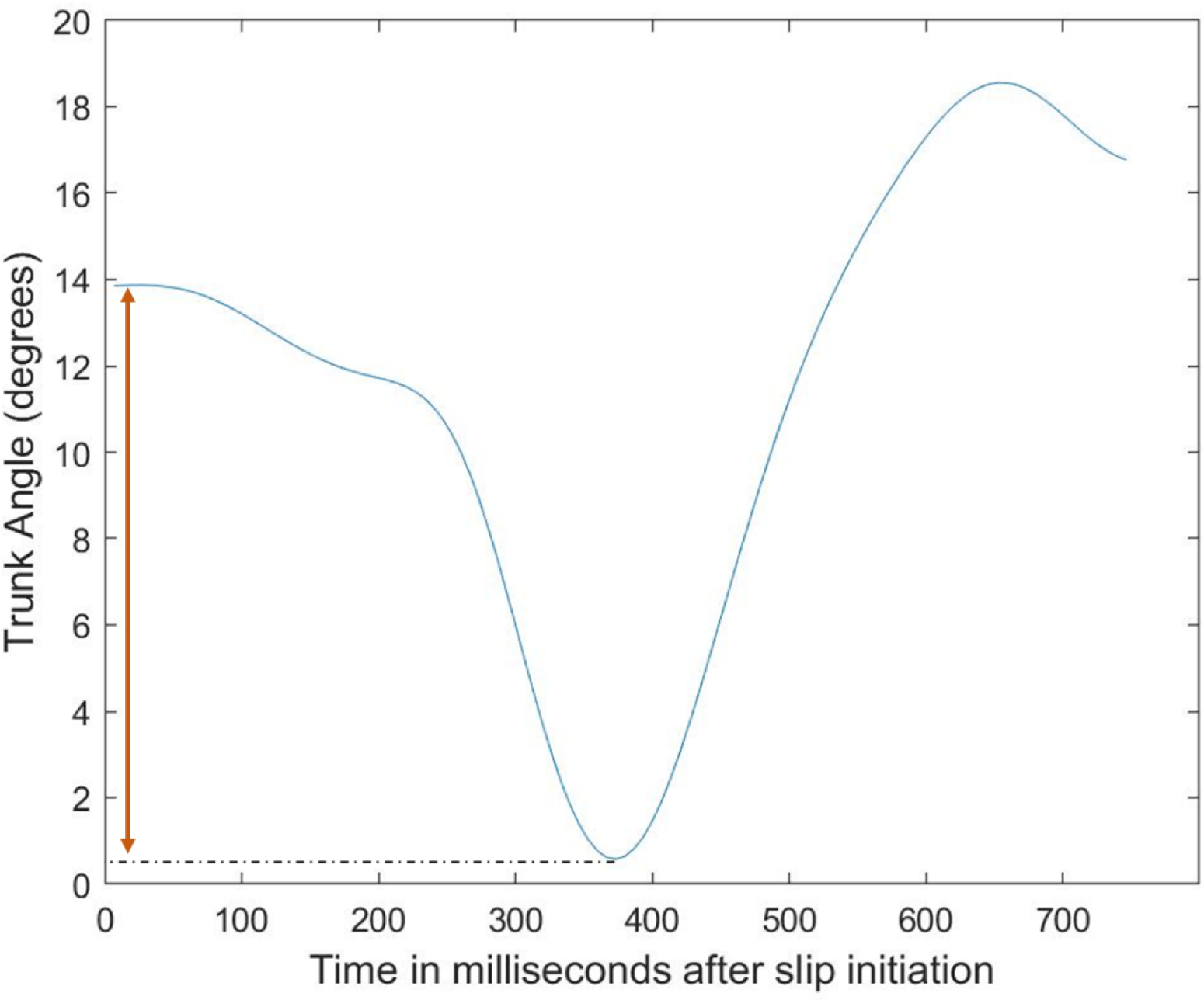
An example from a participant’s sagittal plane trunk behavior over time after a slip initiation. The orange doubled-sided arrow depicts the calculation of excursion. A positive value indicates trunk extension whereas a negative value indicates trunk flexion.

**Figure 2.**
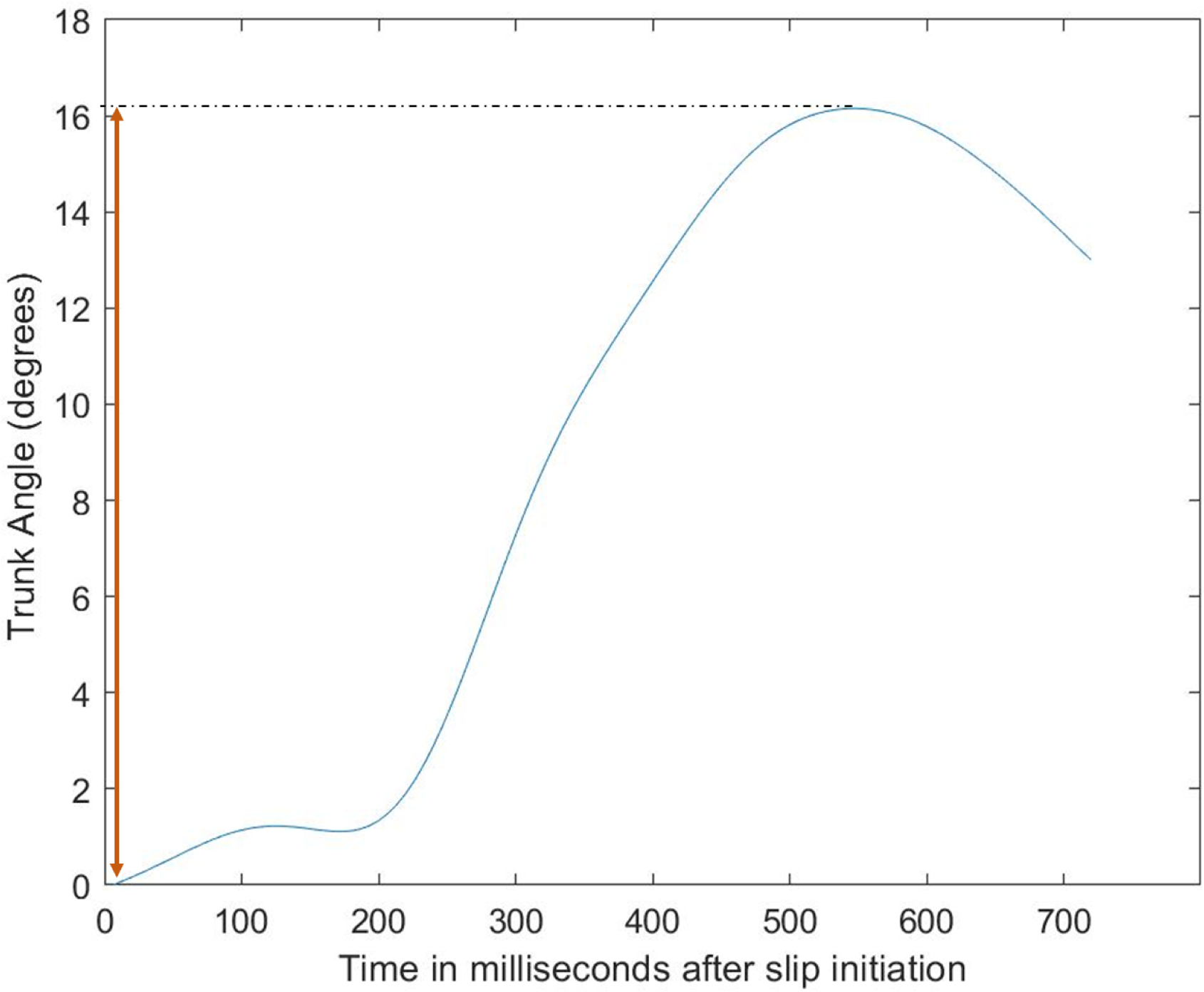
An example from a participant’s frontal plane trunk behavior over time after a slip initiation. The orange doubled-sided arrow depicts the calculation of excursion. A positive value indicates right trunk flexion whereas a negative value indicates left trunk flexion.

### Statistical Analysis

A shapiro-wilk test was conducted to test for normality of the data. A paired t-test was performed to test for differences in peak trunk angles between the sagittal and frontal plane. Statistical analysis was performed using SPSS software (SPSS, Chicago, IL, USA). Significance levels were set at p < 0.05.

## Results

The results depicted below are from the 13 participants who recovered their balance as three individuals fell. Peak trunk angles from the sagittal and frontal plane trunk movements were normally distributed. However, trunk excursion in the sagittal and frontal plane were not normally distributed. As such, trunk excursion comparisons were assessed with a Mann-Whitney U test. The results of the Mann-Whitney U test revealed that there was no significant difference in peak trunk angles between the sagittal and frontal plane (18.7° ± 7.2 vs. 20.4° ± 9.8, respectively, p = 0.73, Fig. 3). The participants’ range for frontal plane trunk right flexion was 3.71 to 32.91 degrees.

**Figure 3.**
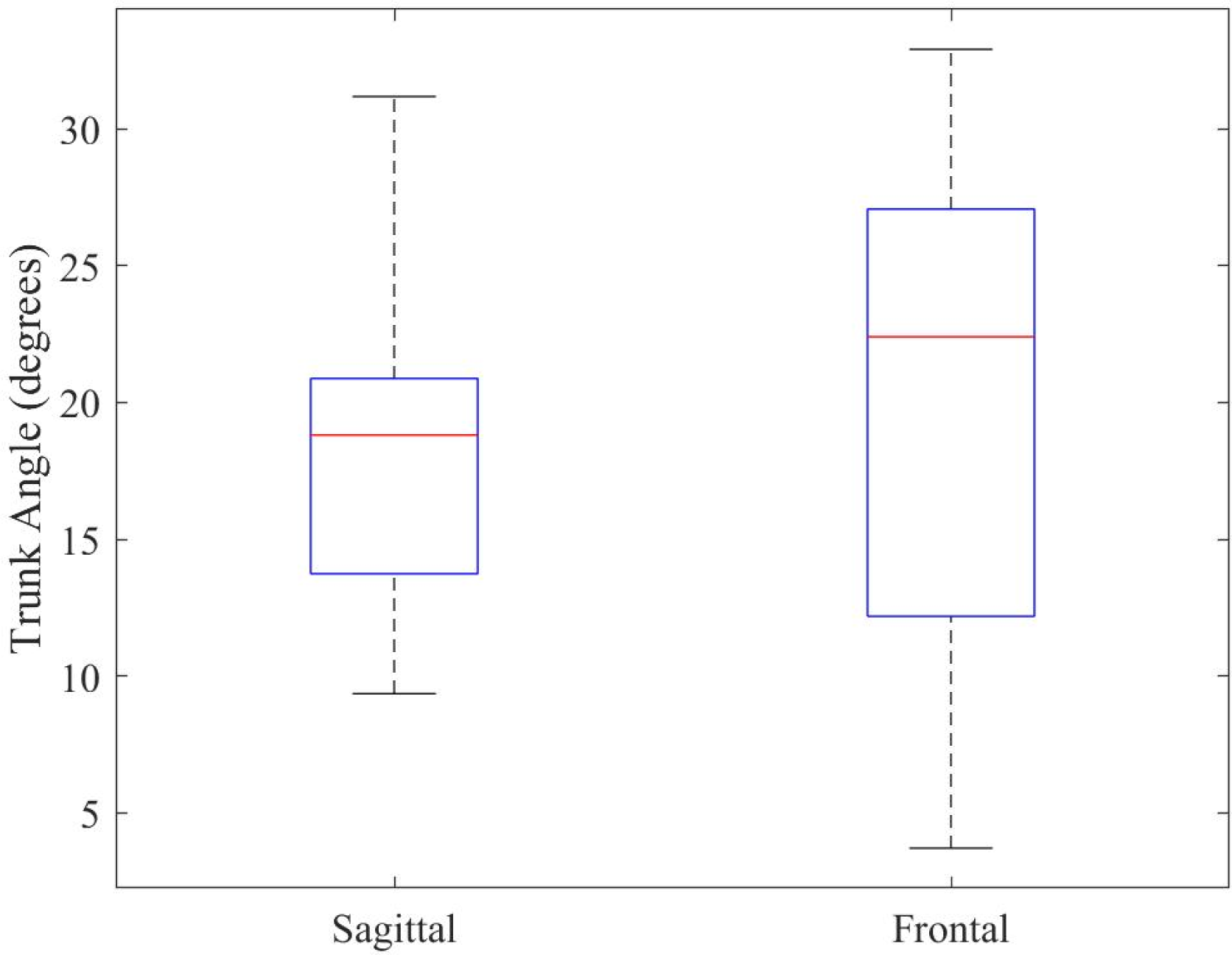
The peak trunk angles of the sagittal and frontal plane after slip initiation (n = 13, p = 0.73). Positive values are trunk extension (sagittal) and right trunk flexion (frontal).

## Discussion

The purpose of the current study was to characterize the trunk motions in the sagittal and frontal planes during a slip incident. There were no significant differences in the peak trunk angles between the sagittal and frontal plane. Surprisingly, the direction of the trunk in the sagittal plane was opposite of what we expected. Overall, these findings suggest that further analyses into frontal plane mechanics of the arms and legs countering trunk motion during a slip are warranted.

The finding of this study differed from other studies that reported a backwards movement within the sagittal plane. Previous studies have reported that a backwards loss of balance occurs after an individual experiences a slip [5,7,9,27–31]. In this study, the initial response of our participants exhibited a trunk flexion response immediately following slip initiation, followed by an overcompensation trunk extension response (Fig. 1). It is possible that the differences in sagittal plane mechanics differ as the methodology to induce a slip differs between studies. Our study utilized mineral oil on the floor as a contaminant, whereas other studies used moveable platforms [30], vinyl covers with detergent [31], and wax paper [24]. Regardless of the differing sagittal mechanics, a focus on the sagittal plane has neglected the importance of analyzing slips in the frontal plane, and this is important as a sideways loss of balance is what can lead to increased risk of hip fractures [14].

There was frontal plane motion of the trunk induced by a slip incident. The initial movement responses of the trunk in the frontal plane were directed towards the side of the slipped foot. In this particular study, all the participants experienced a slip on their right foot and their trunk demonstrated right trunk flexion. To note, individuals that experienced a slip on their left foot exhibited left trunk flexion in our pilot testing suggesting this is a mechanical response and not dependent on dominant or non-dominant limbs. To the best of our knowledge, this study is the first study to address the frontal plane movement of the trunk during a slip and provides support to the claim that slip incidents can cause hip fractures [32].

From a theoretical biomechanical perspective, an overground slip incident during walking would inherently induce frontal plane motion. A slip incident begins with an individual making foot contact onto a slippery surface where the foot intends on accepting weight. During a slip however, the perturbed foot slides anteriorly away from the body while still maintaining weight on the foot as the slip progresses. This weight bearing on the anteriorly shifting perturbed foot would lower the height of the hip on the perturbed side causing the trunk to shift within the frontal plane. A schematic of this theoretical biomechanical movement is demonstrated in Figure 4. This theoretical concept is observable in our participant as shown in Figure 5. As shown in Figure 5, the frontal plane motion of the trunk is exhibited, whereas the sagittal plane excursion is less present.

**Figure 4.**
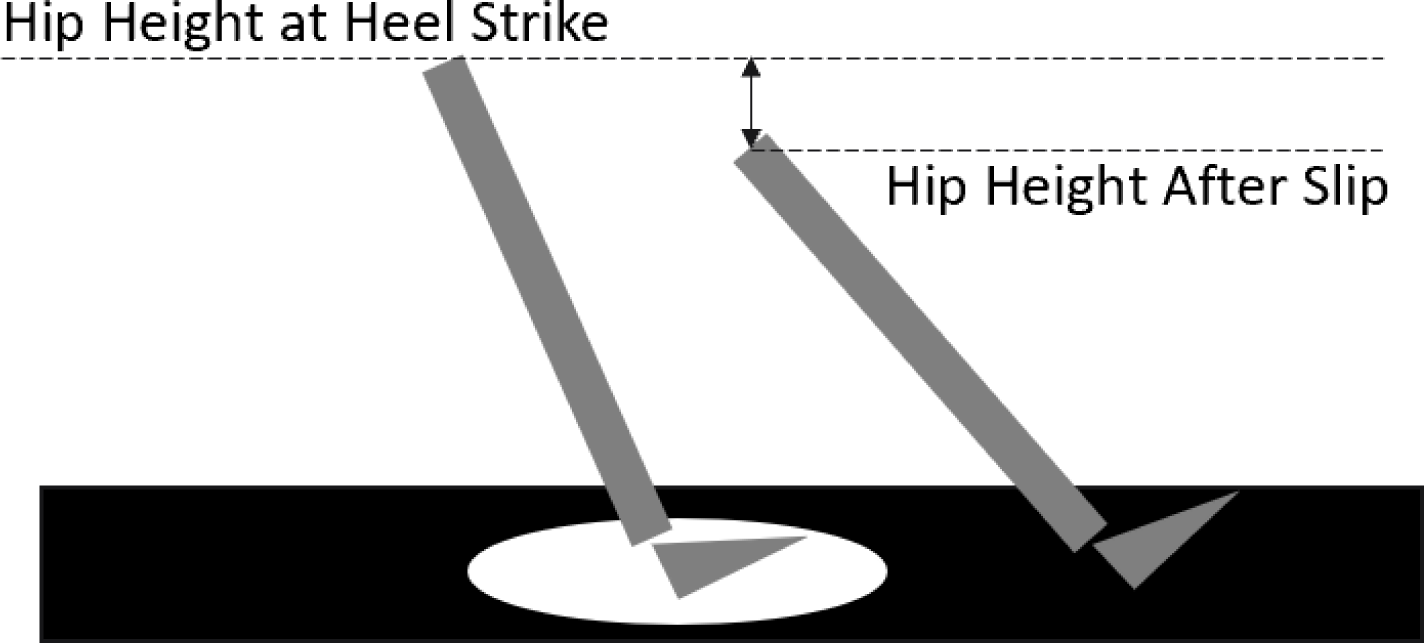
A schematic depicting the theoretical biomechanical inducing of frontal plane motion during a slip incident. The left gray limb is the right leg at heel strike. The white circle represents a slippery substance on the floor. The top horizontal dashed black line represents the hip height at heel strike. The right gray limb represents the right leg during the slip after the foot slides forwards. The bottom horizontal dashed black line represents the hip height after slip initiation. The double-sided vertical arrow represents the hip height difference on the side of the slipped foot between the hip height before and after the slip.

**Figure 5.**
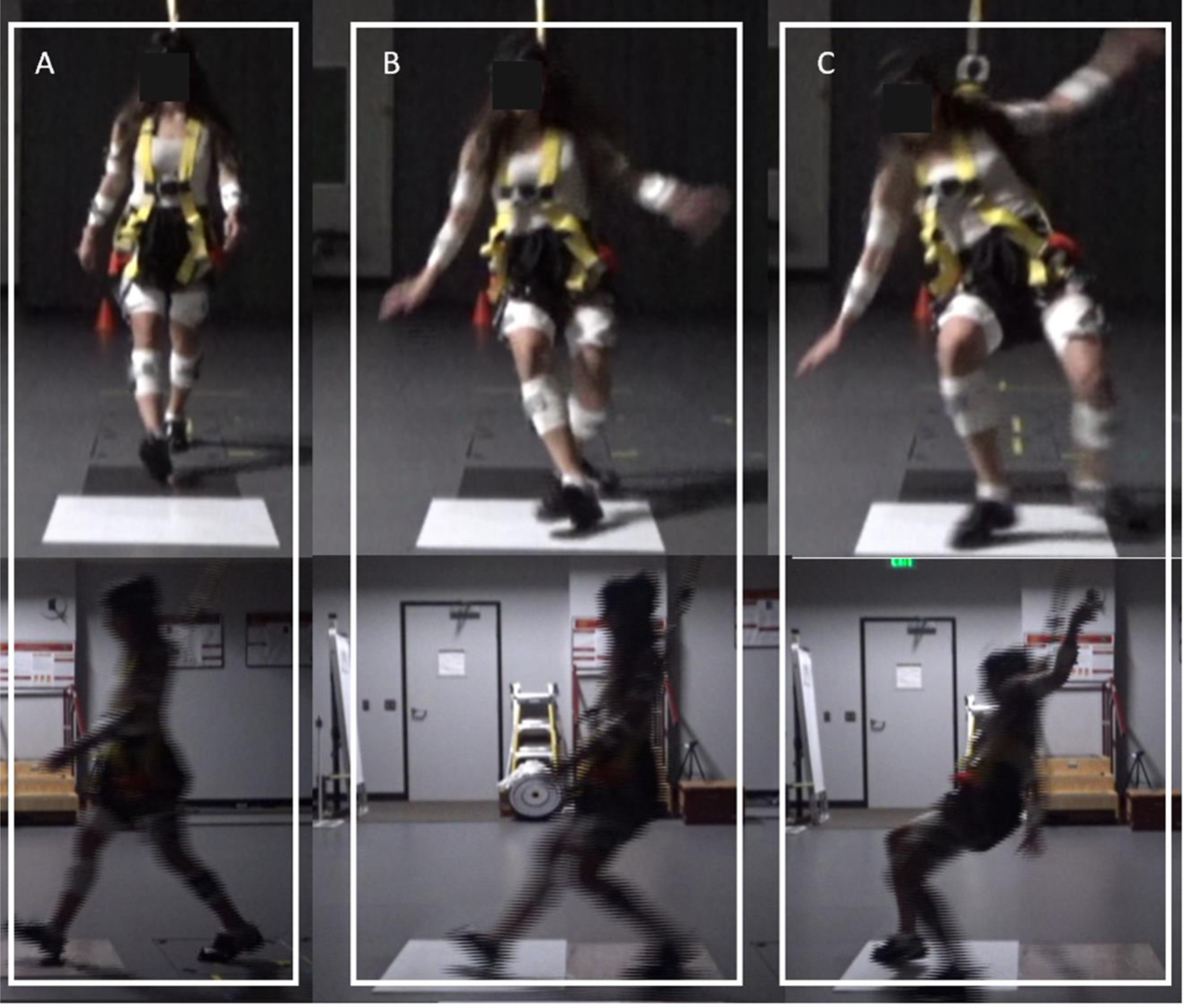
An example of a participant experiencing a slip incident in the laboratory. The photograph depicts three different progressions of the individual undergoing the slip incident. The vertical white box labeled “A” shows a frontal view (top) and sagittal view (bottom) of an individual just prior to heel strike on the slippery surface. The vertical white box labeled “B” shows a frontal view (top) and sagittal view (bottom) of an individual after slip initiation. The vertical white box labeled “C” shows a frontal view (top) and sagittal view (bottom) of an individual towards the end of the slip incident. The frontal plane motion of the trunk is visible in the frontal view, and the sagittal plane rotational behavior of the trunk is quite minimal in this participant.

Frontal plane motion during a slip could be difficult to control. A slip incident results in the body experiencing rapid movements of the extremities and trunk [3,5,8,10,20,31,33]. The head, arm and trunk (HAT) account for approximately 67.8% of total body weight [34]. Having majority of the weight of the body rapidly rotating towards the left or the right would be difficult to control as it carries most of the mass of the body. One study showed that head and trunk velocities were significantly greater in the frontal plane compared to the sagittal plane during a walking treadmill slip [35]. This issue of frontal plane loss of balance becomes more troublesome when applying this concept to older adults. While adults shift postural strategies from an ankle strategy to a hip strategy as they age, it was reported that older adults demonstrated low activation of the rectus abdominus which led to higher trunk movements in a platform perturbation [36]. Furthermore, the legs are focused on regaining balance by restoring the base of support from the antereoposterior disturbance of the slip, and the legs are limited on the frontal plane motion it can contribute to sideways loss of balance. This is likely why the arm movements exhibit frontal plane movement to counter a sideways loss of balance [9,21,37]. It is likely that uncontrolled frontal plane trunk movement during a slip incident is what leads to older adults fracturing their hip as that is the direction which is the most difficult to control [14].

While frontal plane trunk motion is difficult to control, there are ways for individuals to mitigate the severity of frontal plane-induced mechanics from a slip. The arms were shown to act as a counterweight during a sideways platform by having the arms move opposingly to the direction of the perturbation [38]. The counterweight movement of the arms would likely oppose the direction of frontal plane trunk disturbance by minimizing the center of mass trajectory [21,39]. Furthermore, arm abduction during an overground slip incident reduces center of mass dynamics and increases the margin of stability to assist in regaining balance within the frontal plane during a slip incident [21]. Lastly, these frontal plane arm movements are shown to reduce falls 70% during an overground slip [20]. These counter-movements would only be present if a frontal plane trunk disturbance was induced during a slip incident.

There are some considerations that must be addressed when interpreting the results of this study. Firstly, this study recruited and analyzed young and healthy adults so these results should not be generalized to older adults. Further exploration of studies investigating older adults’ mechanics during a slip incident is warranted to generalize findings. Secondly, the sample size of the study was 13 participants which may be considered small, however all participants demonstrated a sideways rotational behavior of the trunk as shown in the photographs. A third consideration is the slip methodology used in this study and others. This study induced a slip during walking using oil on a floor, whereas other studies used treadmills, moveable platforms, wax paper, soapy water & vinyl covers with detergent and do both standing and walking slips. While it is important to note the type of slip methodology for interpretation, we believe the information in this study is still relevant as individuals slip on floors with lower friction values such as restaurants workers stepping on oily surfaces or ice, individuals walking on muddy tiles in outdoor spaces, or individuals stepping on wet bathroom floors.

## Summary

There was a similar amount of frontal plane trunk motion compared to sagittal plane trunk motion providing evidence that the frontal plane mechanics of a slip warrant investigation. The participants in our study exhibited an initial trunk flexion response in the sagittal plane differing from previously reported literature. Frontal plane mechanics observed during a slip warrant further investigation as hip fractures result from individuals falling on their sides.

